# Si-C: method to infer biologically valid super-resolution intact genome structure from single-cell Hi-C data

**DOI:** 10.1101/2020.09.19.304923

**Authors:** Luming Meng, Chenxi Wang, Shi Yi, Qiong Luo

## Abstract

There is a strong demand for the methods that can efficiently reconstruct biologically valid super-resolution intact genome 3D structures from sparse and noise single-cell Hi-C data. Here, we developed Single-Cell Chromosome Conformation Calculator (Si-C) within the Bayesian theory framework and applied this approach to reconstruct intact genome 3D structures from the single-cell Hi-C data of eight G1-phase haploid mouse ES cells. The inferred 100-kb and 10-kb structures consistently reproduce the known conserved features of chromatin organization revealed by independent imaging experiments. The analysis of the 10-kb resolution 3D structures revealed cell-to-cell varying domain structures in individual cells and hyperfine structures in domains, such as loops. An average of 0.2 contact reads per divided bin is sufficient for Si-C to obtain reliable structures. The valid super-resolution structures constructed by Si-C demonstrates the potential for visualizing and investigating interactions between all chromatin loci at genome scale in individual cells.

## Introduction

Chromatin folds into intricate three-dimensional (3D) structure to regulate cellular processes including gene expression^1-4^. Recent researches on enhancer perturbation^5,6^ revealed that many gene regulations are associated with higher-order interactions that involve three or more chromatin loci. A new opinion appears to be becoming popular that enhancer-promoter pairing, both in the sense of the 3D genome and function, is not binary but quantitative^3,7^. The currently reported experiment of super-resolution chromatin tracing confirmed that cooperative higher-order chromatin interactions are widespread in single cells and, moreover, such interactions exhibit substantial cell-to-cell variation^8^. Notably, in mammalian genomes, most enhancers are tens to hundreds of kilobases away from their target promoters^6^. These findings highlight the critical need for super-resolution 3D genome structures of individual cells.

Thanks to advances in genome-wide chromosome conformation capture technology (Hi-C) and decreasing sequencing costs, single-cell Hi-C protocols have been available^9-12^. Computational modeling of genome structure from single-cell Hi-C contact data provides a pivotal avenue to address the critical need^9,13-16^. However, only a small portion of contacts between genomic loci can be probed in a single-cell Hi-C experiment^17^. The extreme sparsity and noisy of contacted information thus poses a huge challenge for determining biologically valid and high-resolution 3D genome structure from single-cell Hi-C data.

A kind of popular methods, known as constraints optimization methods, was widely used to determine 3D structure from Hi-C data^9,10,12,13,18^. These methods describe chromatin fiber as a polymer consisting of beads of the same size and introduce a cost function taking into account input Hi-C data and polymer-physics properties as constraints. Then, an optimization or sampling procedure is performed to minimize the cost function and finally return a structure that satisfies the predefined constraints. In principle, constraints optimization methods implicitly assume that there is a unique 3D structure underlying the single-cell Hi-C data. However, different conformations can satisfy the incomplete Hi-C data and cannot be distinguished by experiments. To overcome the problem, constraints optimization methods repeat the optimization procedure multiple times from different random initial structures to generate structure ensembles. Although such a practice seems plausible, the variability of the structure ensemble cannot reflect the valid structure error bar, since the ensemble lacks a statistical foundation.

To address the issue, Carstens *et al*.^19^ adopted the Bayesian Inferential Structure Determination (ISD)^20^ originally developed for reconstructing structures of proteins from NMR data. The Carstens *et al*. method^19^ builds a posterior probability distribution combining experimental single-cell data with polymer-physics-based model of chromatin fiber. Through using Markov chain Monte Carlo algorithm, a sampling procedure is carried out to produce statistically meaningful structure ensemble that represents the posterior probability. The probabilistic nature of the Carstens *et al*. method makes it efficient for structure determination from sparse and noisy data. However, this method is resource-consuming and thus unsuitable for large-scale genome structure determination.

When we began this study, there was only one published method developed by Laue’s group, named NucDynamics^9^, that has the ability to reconstruct intact genome 3D structures from single-cell Hi-C data of mammalian cells^9,14,21,22^. However, NucDynamics is one of constraints optimization methods which lack a statistical foundation. Here, we developed Single-Cell Chromosome Conformation Calculator (Si-C) within the Bayesian theory framework. Briefly, Si-C resolves a statistical inference problem with a fully data-driven modeling process (for the schematic illustration of Si-C method see Fig. 1). The goal of Si-C is to obtain the 3D genome conformation (**R**) with the highest probability under the given Hi-C contact constraints (**C**) being satisfied. We define the conditional probability distribution *P*(***R***|***C***) over the whole conformation space with Bayesian theorem. After that, the total potential energy of a 3D conformation is defined as *E*(***R***) =*E*_*cont*_ (***R***) + *E*_*phys*_(***R***), where *E*_*cont*_(***R***) = -ln *P*(***R***|***C***) describes the potential energy derived from contact constraints and *E*_*phys*_(***R***) is a harmonic oscillator potential ensuring the connection between consecutive beads, and then an optimization procedure is performed to minimize *E*(***R***), or in other words, to maximize *P*(***R***|***C***). To rapidly achieve the optimized structure at target resolution, we incorporate hierarchical optimization strategy into Si-C framework. For each cell the whole calculation of a given resolution is performed 20 times using the same input contact data, but starting from different random initial structures to finally generate a structural ensemble.

**Fig. 1:**
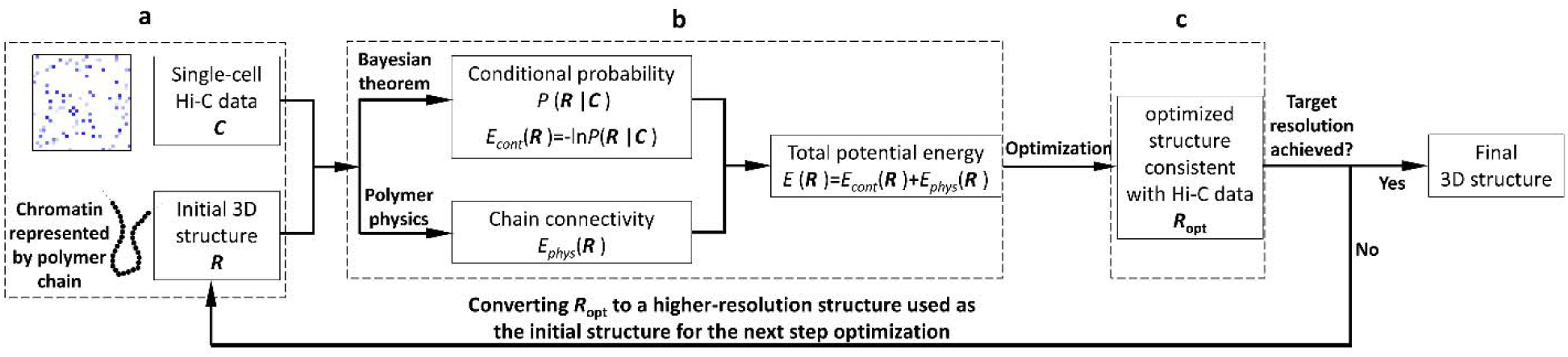
Schematic illustration of Si-C method for inferential structure determination from single-cell Hi-C data. **a** Building models of chromosomes by using polymer chains consisting of contiguous, equally sized beads, and assigning the experimentally observed contacts to the pair of chromatin beads containing the corresponding restriction fragment ends. **b** Defining the total potential energy of a 3D conformation. **c** Performing a structure optimization by using steepest gradient descent algorithm from an initially conformation to minimize *E*(***R***).

Si-C overcomes the limitations of the Carstens *et al*. method and NucDynamics and enables rapid inference of biologically and statistically valid whole-genome structure ensemble at unparalleled high resolution from single-cell Hi-C data of mammalian cell.

## Results

In this study, we applied both of the Si-C and NucDynamics approaches to structure determinations from the single-cell Hi-C data of eight G1-phase haploid mouse embryonic stem (ES) cells^9^(data is downloaded from Gene expression Omnibus or GEO with accession code GSE80280), and compared the validity and performance of Si-C with NucDynamics. Before using the Hi-C data to model 3D structures either by Si-C or NucDynamics, we removed isolated contacts which are not supported by other contacts between the same two regions of 2 Mb because these contact reads have high risk of sequence mapping errors. Details are presented in Methods section and Supplemental Note 1, respectively.

### The percentage of contact restraints that are violated in the 3D model

As a first validation of Si-C, we measured the percentage of experimental contact restraints which are violated in the calculated 3D structures. Contact restraints are violated if the corresponding two beads are separated with distance larger than 2 bead diameters. Fig. 2a shows that the average violation percentages of the Si-C 100-kb resolution structure ensembles of the eight cells are all less than 1%, while those of the published NucDynamics 100-kb structure ensembles of the same eight cells (structures are downloaded from GEO with accession code GSE80280) range from ∼5% to ∼11% except for Cell 8. For further comparison, we ran both of Si-C and NucDynamics to generate the structure ensembles for Cell 1 at various resolutions and presented their average violation percentages in Fig. 2b. It is found that the average violation percentage of the NucDynamics ensemble is consistently higher than that of the Si-C ensemble across all resolutions and it increases to more than 40% when resolution is improved to higher than 100 kb. In contrast, Fig. 2a illustrates that the violation percentage of Si-C decreases to nearly zero when the resolution is improved from 100-kb to 1-kb for the all eight cells. Hence, the Si-C calculated structures are significantly consistent with the experimental contact constraints, and moreover the consistence is stable towards resolution change. These results strongly validate the reliability of the Si-C method.

**Fig. 2:**
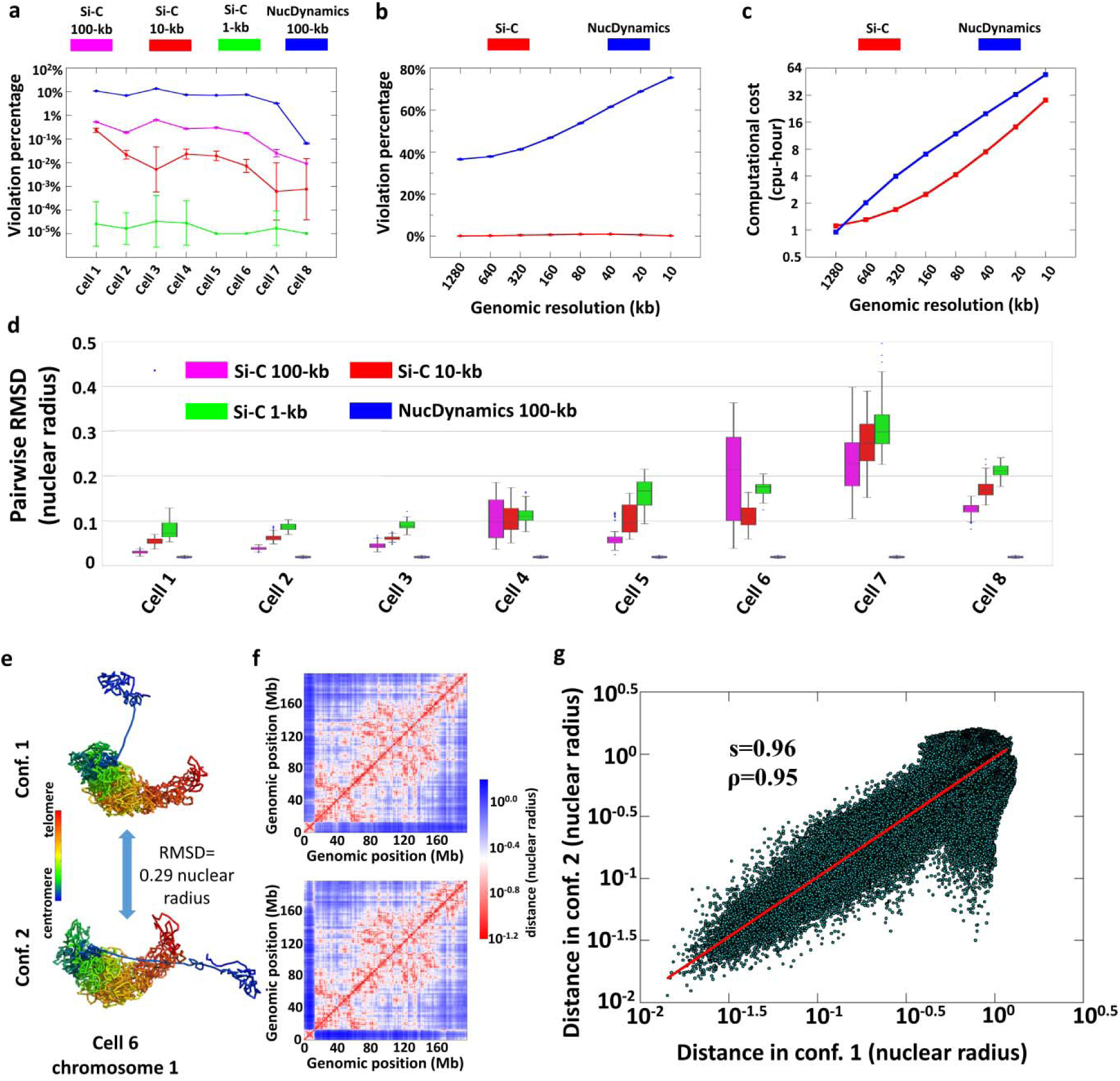
Estimating the validity and performance of Si-C. **a** The average percentages of experimental contact restraints violated in the Si-C ensembles at 100-kb (purple), 10-kb (red), and 1-kb (green) resolutions along with the average violation percentages of the published NucDynamics 100-kb structure ensembles of the same cells (blue) [structures downloaded from GEO with accession code GSE80280]. The distance threshold to decide whether two beads are in contact set to 2 bead diameters. **b** The average percentages of violated contacts in the ensembles of Si-C (red) and NucDynamics (blue) of cell 1 at resolutions ranging from 1280 kb to 10 kb. **c** Comparison of Si-C (red) and NucDynamics (blue) in terms of computational cost for structure reconstruction for cell 1. Details are presented in Supplementary Note 2. Error bars in **a, b** and **c** are the mean ± SD. **d** Boxplot of pairwise RMSDs within the Si-C ensembles at 100-kb (purple), 10-kb (red), and 1-kb (green) resolutions along with pairwise RMSDs within the published NucDynamics ensembles of the same cells (blue) [structures downloaded from GEO with accession code GSE80280]. Median values are shown by black bars. Boxes represent the range from the twenty-fifth to the seventy-fifth percentile. The whiskers represent 1.5 times of the innerquartile range. **e** The 100-kb structures of two conformations of chromosome 1of cell 6 with the RMSD between them of 0.29 nuclear radii. Chromosome regions are colored from blue to red (centromere to telomere). **f** The distance matrices derived from the two 3D structures shown in **e. g** Correlation between the two distance matrices in **f**. The Pearson correlation coefficients (ρ) is 0.95. The red line is power-law fit with scaling exponent (s) equal to 0.96.

### Conformational variability within the calculated structure ensemble

As a second validation, we assessed the degree of conformational variability within the same structure ensemble by pairwise root mean square deviation (RMSD)^23^. The details of RMSD calculation are described in Supplementary Note 3. For cell 1, 2, and 3, Fig. 2d shows that the variabilities in the Si-C ensembles at three resolutions (median RMSDs <0.1 nuclear radii) are comparable to the published NucDynamics 100-kb structure ensembles^9^. However, the Si-C method seems to produce very heterogeneous ensembles for cell 6 and 7 in terms of the values of RMSDs. Specially, the median RMSD of the Si-C 100-kb structure ensemble of cell 6 is more than 0.2 nuclear radii.

It should be noted that, for the case of reconstructing 3D structure of chromosome represented by polymer chain consisting of numerous beads, high RMSD between two conformations might be caused by the flexibility of the polymer chain, the inconsistence in chromatin folding between two conformations, or both of them. To investigate the rationales behind high RMSDs within the Si-C ensemble, we selected two conformations of chromosome 1 from the 100-kb structure ensemble of cell 6 (the RMSD between them is as high as 0.29 nuclear radii) and displayed their 3D structures in Fig. 2e. A quick glance shows that both of the two structures can be viewed to consist of two parts which are connected by a flexible chain and the relative locations of the two parts in the two conformations are apparently different. To quantitatively assess the difference in the chromatin folding between the two conformations, we computed a distance matrix for each conformation (see Fig. 2f; details of distance matrix calculation are presented in Supplementary Note 4). The two distance matrices show high correlation with each other, with Pearson correlation coefficient of 0.96 (Fig. 2g). The high correlation indicates that the flexibility of the polymer chain predominantly contributes to the high RMSD between the two conformations. In other words, the results demonstrate that the Si-C method enables to produce highly consistent structures in terms of chromatin folding.

On the other hand, the Si-C ensembles at different resolutions of the same cell show similar structural variability. For instance, the median RMSDs of the Si-C 100-kb, 10-kb, and 1-kb resolution structure ensembles are (0.02, 0.04, and 0.06 nuclear radii) for cell 1 and (0.21, 0.12, and 0.18 nuclear radii) for cell 6, respectively. The differences between median RMSDs of the Si-C ensembles at different resolutions of the same cell are small, indicating the validity of the Si-C approach.

### Agreement between the experimental and back-calculated data

To quantitatively evaluate the agreement between 3D models and Hi-C maps, we translated 3D models into distance matrices (details are presented in Supplementary Note 4). For example, we constructed the distance matrices for the 10-Mb genomic regions (Chr1: 30 Mb to 40 Mb) based on the Si-C 10-kb structures of the eight individual cells (see Fig. 3a) and averaged them across the eight cells to obtain an average distance matrix (see Fig. 3c) that can be compared to the population Hi-C contact frequency matrix^24^ (Fig. 3b; populated Hi-C data downloaded from GEO with accession code GSE35156). The spatial distance in Fig. 3c displays high correlation with the Hi-C contact frequency in Fig. 3b, with Pearson correlation coefficient of -0.92 (see Fig. 3d). Furthermore, the average distance matrix (see Fig. 3c) shows domain structures similar to Topological-associated domains (TAD) observed in the population Hi-C matrix (see Fig. 3b). We refer to these domain structures as TAD-like structures and identify their boundaries using separation score (details of separation score calculation are presented in Supplementary Note 5). As shown in Fig. 3f, the positions corresponding to the separation score peaks are considered to be boundaries of TAD-like domains. Fig. 3e shows the positions of TAD boundaries in the population Hi-C map which are identified by the TopDom algorithm^25^ (details for TAD boundary identification are presented in Supplementary Note 5). The results clearly show that most of boundaries of TAD-like structures in Fig. 3f align with the positions of TAD boundaries in Fig 3e. These findings demonstrate the significant similarity between the calculated average distance matrix and the population Hi-C contact-frequency matrix, strongly validating the 10-kb resolution structures generated by Si-C.

**Fig. 3:**
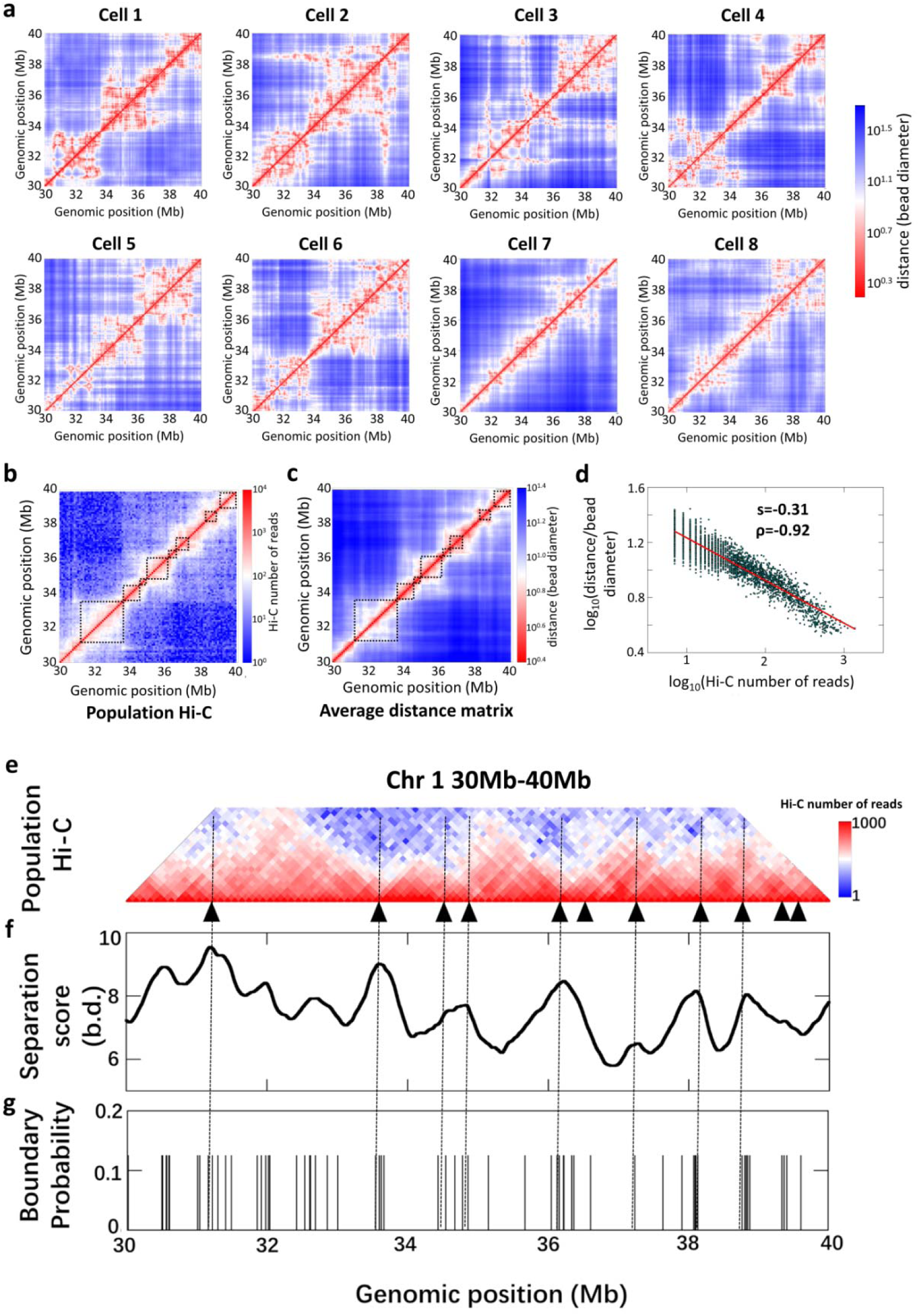
Si-C calculated structures allow identification of TADs in the population Hi-C contact frequency matrix. **a** The 10-kb resolution distance matrices for the 10-Mb genomic regions (Chr1: 30 Mb to 40 Mb) of the eight individual mouse ES cells calculated from the corresponding Si-C 10-kb resolution structures. **b** The 100-kb resolution population Hi-C contact frequency matrix of mouse ES cells for the same 10-Mb genomic region (Chr1: 30 Mb to 40 Mb) [data downloaded from GEO with accession code GSE35156]. TADs identified previously are shown in dark boxes. **c** The 100-kb resolution average distance matrix derived from the eight 10-kb resolution distance matrices shown in Fig. 3a. TAD-like regions identified by separation score are shown in dark boxes. **d** Correlation between the spatial distance and Hi-C contact frequency shown in Fig. 3c and 3b. The Pearson correlation coefficient (ρ) is - 0.92. The red line is power-law fit with scaling exponent equal to -0.3. **e** Enlarged view of Fig. 3b with the positions of TAD boundaries labeled by black triangles. **f** Separation score for each genomic position in Fig. 3c. **g** Probability for each genomic position to appear as a single-cell domain boundary. Single-cell domain boundaries are identified by computing separation score for each genomic position of each distance matrices shown in Fig. 3a.

A closer inspection of Fig. 3a immediately shows that domain structure is also a fundamental hallmark of genome organization of individual cells, and the sizes and locations of these domains display substantial cell-to-cell variations. These findings are consistent with previous super-resolution imaging experiment^8^. To carry out a quantitative comparison with the observation of the super-resolution imaging experiment^8^, we also identified domain boundaries in the eight matrices shown in Fig. 3a and measured the probability for each genomic position appearing as a single-cell domain boundary throughout the eight matrices. Notably, the 10-Mb region is represented by 1000 beads at the 10 kb resolution among which only ∽6% of beads appear as a single-cell boundary in any of 8 cells with nearly the same probability of 0.125 (Fig. 3g). Whilst the number of cells is small, the statistical results presented in Fig. 3g demonstrate two pronounced trends. First, the number of positions which appear as a single-cell domain boundary is evidently higher than the total number of established TAD boundaries in the populated Hi-C matrix (Fig. 3e). Second, the regions where the single-cell boundaries are enriched are well aligned with the positions of TAD boundaries. These trends are in agreement with the observations of the super-resolution chromatin tracing experiment that single-cell domain boundaries occur with nonzero probability at all genomic positions but preferentially at the positions which are identified as TAD boundaries in the Hi-C contact frequency map^8^. The agreement provides another support for the validities of the 10-kb resolution Si-C structures.

### Conserved 3D whole-genome architecture in all cells

Several conserved features of 3D genome architecture are consistently observed in the Si-C structures at different resolutions for all 8 G1-phase haploid ES cells. Fig. 4a and 4B display the features of cell 1, while those of the other seven cells are shown in Supplementary Fig.1. First, Fig. 4a shows individual chromosomes occupy distinct territories with a degree of chromosome intermingling (Supplementary Fig.3, calculation details are presented in Supplementary Note 7), and meanwhile centromeres and telomeres localize at opposite sides of the nucleus (Supplementary Fig.2). Second, each territory is divided into euchromatic and heterochromatic regions, known as A and B compartments (Fig. 4c), which is consistent with imaging experiments^26^. (Details of compartment identification are presented in Supplementary Note 6). Third, all chromosomes stack together to give a sphere consisting of two shells and a core: an outer B compartment shell, an inner A compartment shell, and an internal B compartment hollow core (at the 100-kb resolution) or seemingly solid core (at the 10-kb or 1-kb resolutions) (see Fig. 4b). Mapping various experimental data, such as lamina-associated domain^27^ (LAD), CpG density, and DNA replication timing^28^ (data obtained from ArrayExpress with accession code E-MTAB-3506), onto the 3D models shows similar architectures (Fig. 4b), demonstrating that active segments are predominately located in the inner A compartment shell while inactive segments mostly distribute in the outer or internal B compartment regions. These conserved features of intact genome architecture showing in the Si-C structures are consistent with independent observations^29-32^, demonstrating that Si-C enable to infer biologically reliable 3D genome structures from single-cell Hi-C data.

**Fig. 4:**
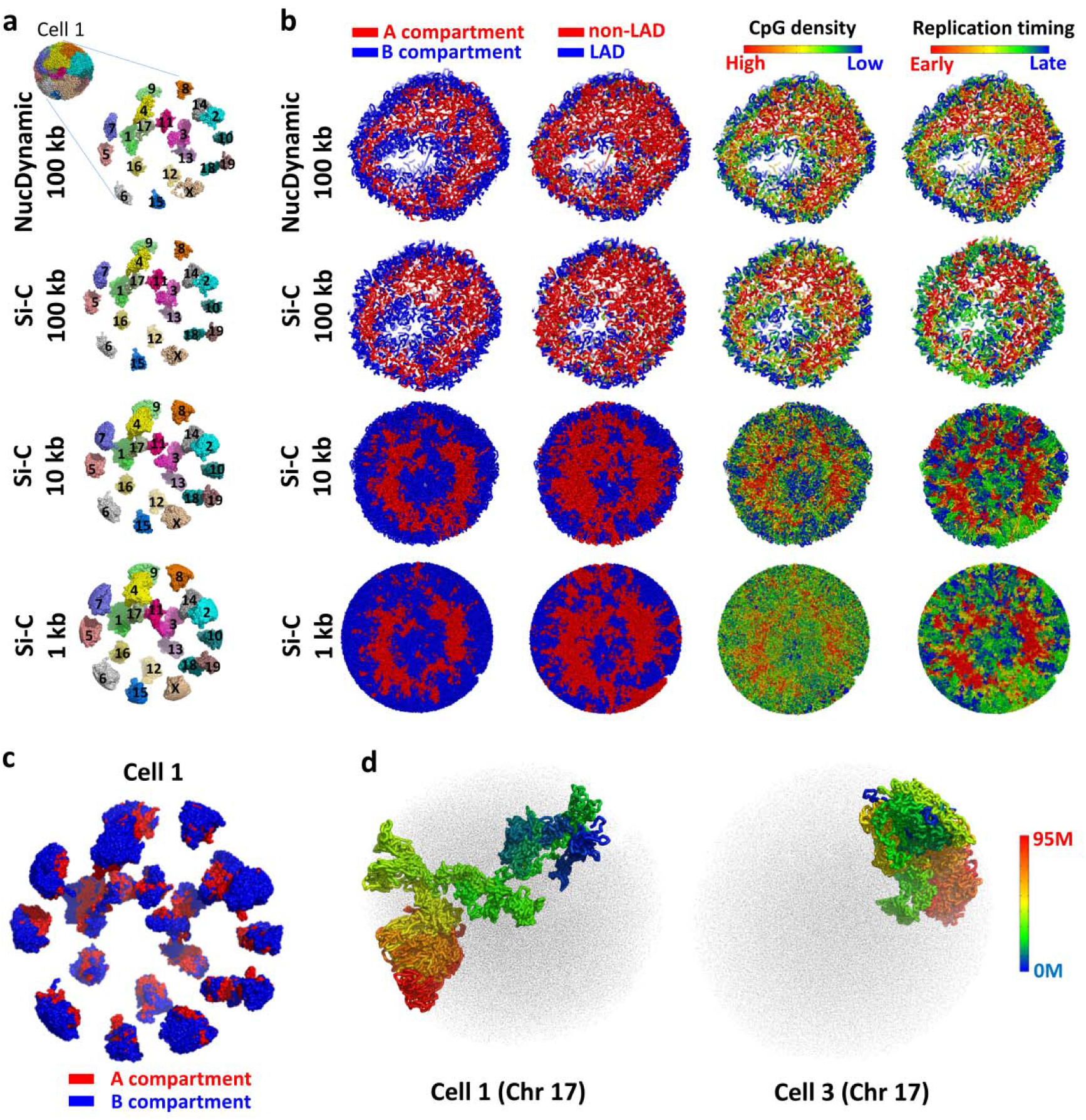
Large-scale 3D structure of the genome. **a** The Si-C calculated intact genome 3D structures of cell 1 at 100-kb, 10-kb, and 1-kb resolutions with expanded view of the separated chromosome territories along with the published NucDynamics 100-kb resolution structure of cell 1. The 100-kb resolution structures of NucDynamics are downloaded from GEO with accession code GSE80280. **b** Cross-sections of intact genome 3D structures of cell 1, colored according to whether the sequence is in the A (red) or B (blue) compartment (first column); whether the sequence is part of a lamina associated domain (LAD) (blue) or not (red) (second column); the CpG density from red to blue (high to low) (third column); the replication time in the DNA duplication process from red to blue (early to late) (fourth column). **c** The Si-C calculated 10-kb resolution intact genome 3D structure of cell 1 with expanded view of the spatial distribution of the A (red) and B (blue) compartments. **d** The Si-C calculated 10-kb resolution structures of chromosome 17 of cell 1 and 3 colored from red to blue (centromere to telomere) with structures of other chromosomes shown as black dots.

Fig. 4d displays another example to illustrate independent evaluation of the validity of Si-C. Although the features of 3D genome architectures shown in Fig. 4a, 4b, and 4c are conserved in all eight cells, the individual chromosome structure varies remarkably from cell to cell (Fig.4d). The cell-to-cell variation is agreement with the observation of previous single-cell experiments^8,9,16^.

### Unparalleled details of chromatin folding provided by Si-C models

The Si-C method enables to rapidly reconstruct whole-genome structures at 100 kb, 10 kb, and even 1 kb resolutions from the single-cell Hi-C data of mouse ES cells^9^. The validity of the Si-C models of 10 kb and 100 kb resolutions are supported by various measurements as mentioned above, while the NucDynamics models of resolutions higher than 100 kb are not credible because the percentages of experimental contact restraints that are satisfied in such models are less than 60%. Here, we showed details of chromatin folding based on the Si-C 10-kb 3D structures and compared them with that of 100-kb structures. For instance, Fig. 5 shows the 10-kb resolution 3D structures of four different regions, each of which includes two neighboring domains. It can be seen that chromatin forms distinct spatially segregated globular domain structures (Fig. 5b), which is supported by the super-resolution imaging experiment^8^. Notably, domains are linked by highly extended chromatin chains. To quantitatively estimate the degrees of compaction of domains and linking chains, we computed the gyration radius (*R*_*g*_) which is defined as the root-mean square distance of bead positions in each 200 kb fragment from the centroid of the 200-kb fragment (details of calculating *R*_*g*_ are presented in Supplementary Note 8), and we assigned the value of *R*_*g*_ to the midpoint bead of the 200-kb fragment. After taking every bead in the four 10-Mb regions as a midpoint of a 200-kb fragment, we calculated the *R*_*g*_ for a total number of 4000 fragments and showed the results in the bottom of Fig. 5a. The numerical distributions of gyration radius for the four regions consistently demonstrate that linking chains are highly extended and align with domain boundaries. It is noteworthy that domains also embody extended fragments whose values of *R*_*g*_ are very close to those of linking chains, implying that domain is not a uniformly compacted structure and then there might be detail structures. Indeed, loop and hairpin-like hyperfine structures are observed in domains (see Fig. 5c and 5d). For comparison, 100-kb resolution models of the same regions are also displayed. It can be seen that loops or hairpin-like structures are obscured or even cannot be visualized in the 100-kb resolution structures. Obviously, analysis of the 10-kb resolution structures provides unparalleled details of chromatin folding.

**Fig. 5:**
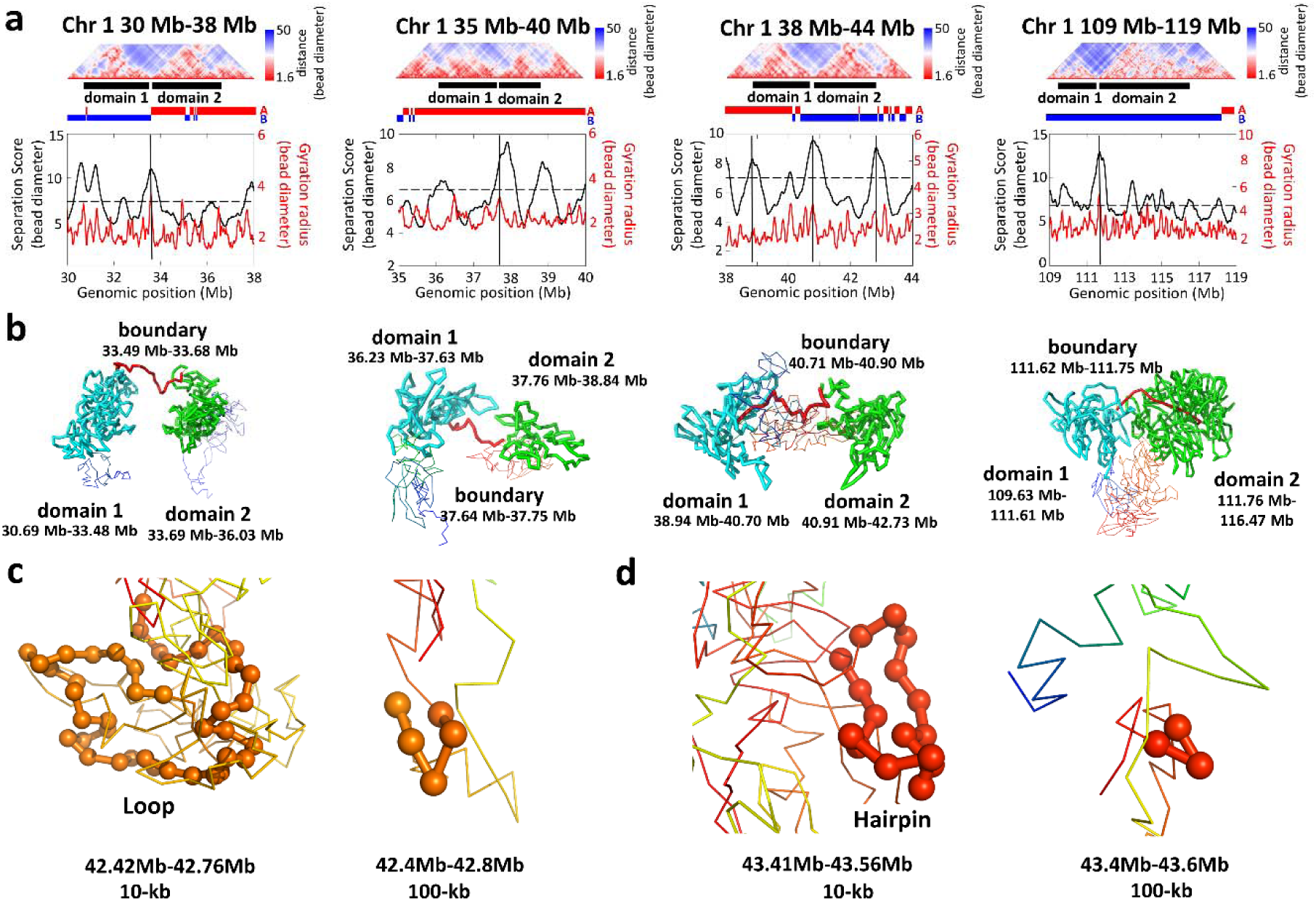
Small-scale 3D structure at 10-kb resolution. **a** Top: heatmap plot of distance matrices derived from the Si-C 10-kb structure of cell 1 for four genomic regions. In each region, two neighboring domains that are identified by separation score are indicated by black lines. Bottom: distributions of the separation score and gyration radius (*R*_*g*_). **b** 3D structures of the genome regions associated with the two neighboring domains shown in Fig. 5a. Structures corresponding to boundary, domain 1 and domain 2 regions are shown in red, cyan, and green, respectively. **c** and **d** Loop and hairpin-like hyperfine structures visualized by beads of 10-kb size (left) and the 3D structures of the same regions visualized by beads of 100-kb size for comparison (right).

### High computational efficiency of Si-C

Last, we compared the execution speeds of Si-C and NucDynamics for reconstructing intact genome structures from the same single-cell Hi-C data at different resolutions (see Fig. 2c; details of computing resources are presented Supplementary Note 2). The comparison shows that, except for 1280 kb resolution, the execution speed of Si-C is twice as fast as that of NucDynamics. Specifically, it costs 28 cpu-hour for Si-C to infer a 10 kb resolution intact genome structure for cell 1, while it is 54.3 cpu-hour for NucDynamics.

## Discussion

In this work, we proposed a Bayesian probability based method, Si-C, to infer the 3D genome structure (**R**) with the highest probability in the light of sparse and noisy single-cell Hi-C data (**C**). Compared with the NucDynamics approach which is the only published method that can determine whole genome structures from single-cell Hi-C data of mammalian cells, Si-C substantially outperformed NucDynamics at rapidly determining biologically and statistically valid super-resolution whole-genome 3D structure. The superiority of Si-C can be attributed to the follows. First, Si-C directly describes the single-cell Hi-C contact restraints using an inverted-S shaped probability function of the distance between the contacted locus pair, instead of translating the binary contact into an estimated distance. For locus pairs without contact reads, an S shaped probability function is used to describe the cases where two loci are spatially close to each other but the proximity is not detected in the Hi-C experiment. The statistical foundation makes Si-C efficient for structure determination from extreme sparse and noisy data. Second, Si-C adopts the steepest gradient descent algorithm to maximize the conditional probability *P*(***R***|***C***), which allows us to rapid inference of an intact genome structure from single-cell Hi-C data of mammalian cells at resolution varying from 100 kb to 1 kb.

For a given single-cell Hi-C dataset, when the resolution of calculated structure improves, the number of beads that are not constrained by experimental data increases, and then the overall precision of inferred structure decrease. Naturally, a question is raised: at least how many contact reads are required for the Si-C method to achieve a reliable 3D structure. Since the median of pairwise RMSDs within the calculated structure ensemble fluctuates along with the change in resolution, we addressed the question through analyzing the relationship between resolution and the median of pairwise RMSDs within the Si-C inferred structure ensemble. We assumed that an inferred ensemble is reliable if its median value of pairwise RMSDs is less than 4 bead diameters. Fig. 6 shows the plot of the median value of pairwise RMSDs within structure ensembles of cell 1 against resolution. According to the threshold of 4 bead diameters, Fig. 6 demonstrates that the highest attainable resolution is 4 kb. At such resolution, the whole genome of cell 1 is divided into 658,453 beads, while the number of Hi-C contact read is 110,623 (each bead corresponds to an average of 0.168 contact reads). Therefore, to obtain reliable 3D genome structure, we suggest that the resolution should be selected such that the amount of contact reads per chromatin bead should be no less than 0.2.

**Fig. 6:**
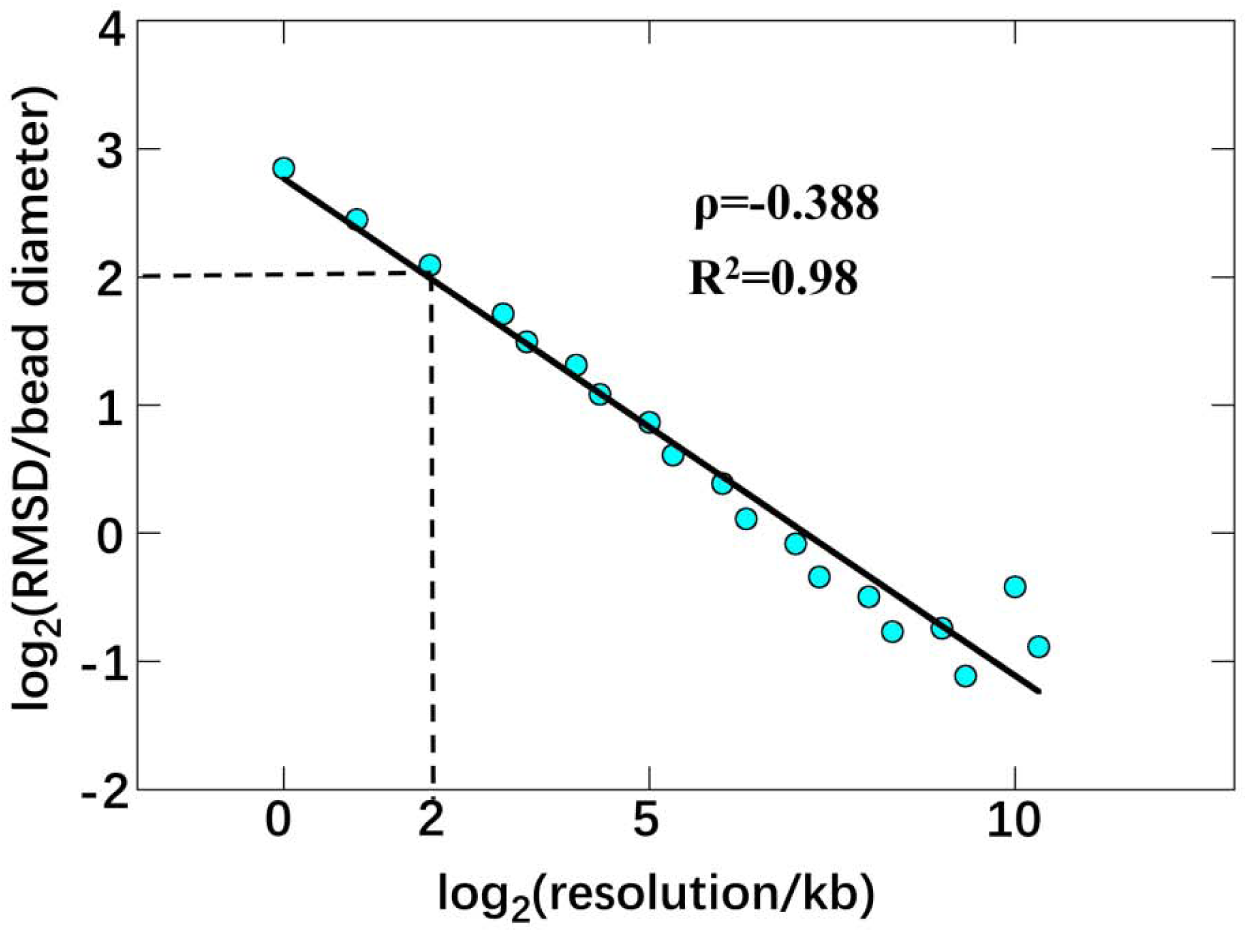
Estimating how many contact reads per chromatin bead are required for Si-C to achieve reliable 3D structure. Log-scale plot of median RMSD within the structure ensemble of cell 1 against resolution. The threshold of the median value of pairwise RMSDs for a reliable calculated structure ensemble set to be 4 bead diameters, indicated by the dash line.

In conclusion, our work shows that Si-C is a biologically and statistically viable method allowing us to rapidly infer a reliable 3D structure at super-resolution which provides unprecedented details on chromatin folding.

## Methods

### Bead-on-a-string representation of chromosome

Each chromosome is represented as a polymer chain consisting of contiguous, equally sized beads with diameter of *a*. The connectivity between adjacent beads (*i,i*+1) is described by the harmonic oscillator potential:

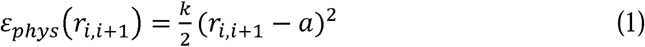

where the force constant *k* is set to 100/*a*^2^ and *r*_*i,i*+1_ is the Euclidean distance between the *i*^*th*^ and *i*+1^*th*^ beads. It should be noted that the diameter *a* is used as unit length in our method.

### Derivation of conditional probability

The basic aim of Si-C is to find out the 3D genome structure (***R***) with the highest probability in the light of the contact restraints (***C***) derived from single-cell Hi-C contact data. Formally, such structure determination can be solved by the probability distribution *P*(***R***|***C***) defined over the complete conformation space. In our approach, the conditional probability *P*(***R***|***C***), also called posterior probability, is defined as:

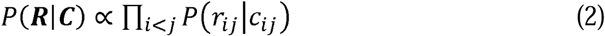

where *r*_*ij*_ is the Euclidean distance between pair of the i^*th*^ and j^*th*^ beads, denoted as beads (*i,j*), and *c*_*ij*_ is the count of detected contact reads between beads (*i,j*) in the single-cell Hi-C matrix. According to the Bayesian theorem, the posterior probability *P*(*r*_*ij*_|*c*_*ij*_) can be factorized into two components:

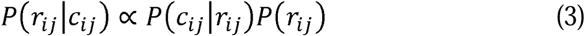

where *P*(*c*_*ij*_|*r*_*ij*_) is the likelihood and *P*(*r*_*ij*_) the prior probability.

The likelihood *P*(*c*_*ij*_|*r*_*ij*_) quantifies the consistence between the contact data and the structural model. It is the probability of observing *c*_*ij*_ contacts between beads (*i,j*) when they are spatially separated by the distance of *r*_*ij*_. Based on the value of *c*_*ij*_, all beads pairs can be divided into two groups. In one group where the value of *c*_*ij*_ is not equal to zero *c*_*ij*_ ≠ 0), the beads (*i,j*) is restrained in close spatial proximity. The *P*(*c*_*ij*_≠ 0 |*r*_*ij*_) would decrease with the increasing of *r*_*ij*_. Therefore, when *r*_*ij*_ is over than the distance threshold *r*_0_, the corresponding *P*(*c*_*ij*_≠ 0 |*r*_*ij*_ > *r*_0_) would be zero. Here, we use an inverted-S shaped probability function to transform the contact constraints derived from single-cell Hi-C data into a function of the distance. The inverted-S shaped probability function is

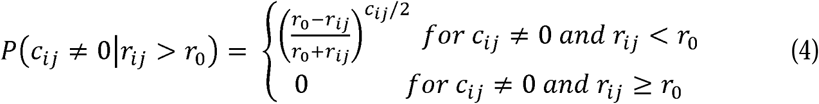

where the distance threshold *r*_*0*_ is set to 2*a*.

In the other group where the value of *c*_*ij*_ is equal to zero (*c*_*ij*_ = 0), the absence of contact read between the beads (*i,j*) cannot be interpreted only as far separation between the beads (*i,j*). There are two distinct rationales behind the absence. The first one is that the absence is indeed because the beads (*i,j*) are too far apart, namely *r*_*ij*_ > *r*_0_. Therefore, *P*(*c*_*ij*_= 0 | *r*_*ij*_ > *r*_0_)approximately equals to 1. The second case is that the beads (*i,j*) are closely located but not detected. This situation is more likely to occur for the pairs in which beads are separated by a large distance but still within the threshold *r*_0_. For this situation, the likelihood *P*(*c*_*ij*_= 0 | *r*_*ij*_ < *r*_0_)is modeled by an S-shaped probability function:

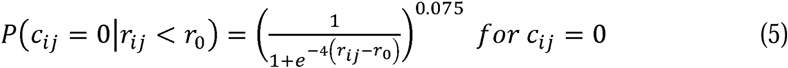

The power coefficient 0.075 is an arbitrary value used in this formula to increase the probability.

The prior probability *P*(*r*_*ij*_) describes the probability of the distance *r*_*ij*_ between beads (*i,j*) based on our prior knowledge. Assuming chromatin beads are uniformly distributed in nuclear space, the prior probability *P*(*r*_*ij*_) is proportional to the surface area of the sphere with radius of *r*_*ij*_. Therefore, the *P*(*r*_*ij*_) is defined as:

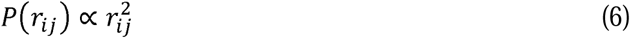

Finally, the posterior probability *P*(*r*_*ij*_|*C*_*ij*_) is adopted as follows:

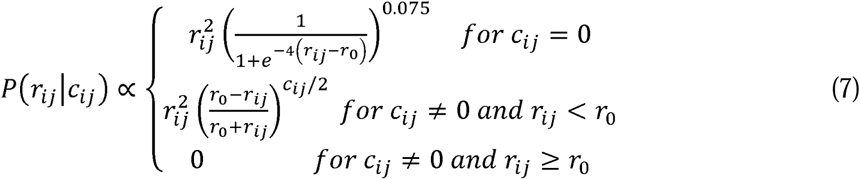

### Total potential energy of a 3D conformation

We define the total potential energy of a 3D conformation with two terms, one of which describes the contact restraints and the second is the above mentioned harmonic oscillator potential introduced to ensure the connectivity of the chromosome backbone. The negative logarithm of the posterior probability, namely −*lnP*(*r*_*ij*_|*C*_*ij*_), is analogous to a physical energy, denoted as *ε*_*cont*_ (*r*_*ij*_,*c*_*ij*_).The *ε*_*cont*_ (*r*_*ij*_,*c*_*ij*_) is used to describe the potential energy originated from the contact restraints. Combining *ε*_*cont*_ (*r*_*ij*_,*c*_*ij*_) with the harmonic oscillator potential *ε*_*phys*_(*r*_*i,i*+1_), the total potential energy *E*(***R***) of a 3D conformation is calculated according to the formula below:

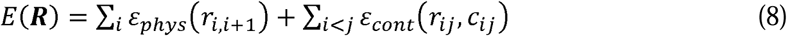

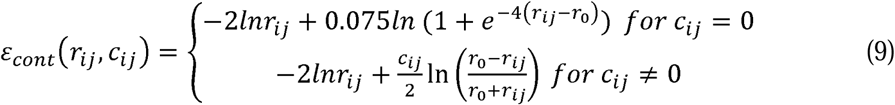

It should be noted that, according to eq. (7), when *r*_*ij*_ increases towards or beyond the distance threshold *r*_0_ for the case of *c*_*ij*_ ≠ 0, the posterior probability decrease to 0 that makes −*lnP*(*r*_*ij*_|*C*_*ij*_) nonsense. To address this issue, we employ Taylor expansion to formulate the energy *ε*_*ij*_ (*r*_*ij*_,*> 0.995r*0*cij≠ 0* as the following:

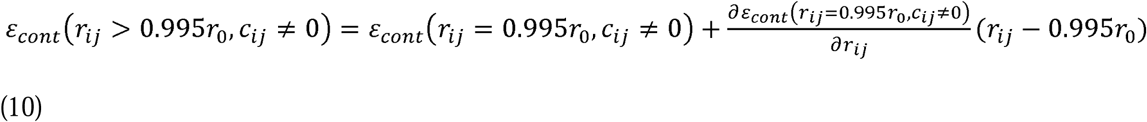

For the bead-on-a-string model of a whole-genome structure, the number of bead pairs belonging to the group of *c*_*ij*_ =0 is huge. To reduce the computational cost, we set the *ε*_*cont*_ (*r*_*ij*_ > 3*a, c*_*ij*_ = 0) equal to the *ε*_*ij*_ (*r*_*ij*_= 3*a, c*_*ij*_ = 0) during the calculation.

### Maximization of conditional probability

3D genome structure optimization is performed by using steepest gradient descent algorithm from an initially random conformation to maximize the conditional probability *P*(***R***|***C***) and output the corresponding structure that is compatible with the experimental contact restraints. In each optimization step, the force that acts on the *i*^th^ bead, denoted as 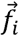, is firstly calculated as the negative derivative of potential energy on the bead coordinate,

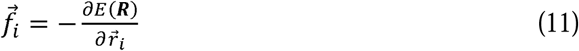

Then the displacement 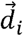 is set to be proportional to the calculated force,

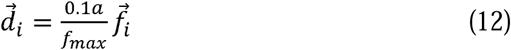

in which *a* is the bead diameter and *f*_*max*_ is the maximum force among the forces on every bead. In the first 2000 optimization steps, the modeled structure is shrunk once per 10 steps. The structure shrinking is performed by multiplying the coordinates of all beads with a factor 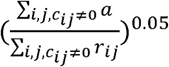, in which *r*_*ij*_ is the distance between the *i*^th^ and *j*^th^ beads and *a* is the bead diameter. After the first 2000 optimization steps, another 8000 optimization steps without structure shrinking are carried out to output the finally optimized structure.

### Hierarchical optimization strategy

The optimized structure of low resolution is used to generate initial structure of high resolution for another optimization procedure to obtain high resolution optimized structure. There are two steps for deriving a high resolution initial structure from the optimized structure of low resolution. First, each bead of the optimized structure of low resolution is evenly divided into two small beads. Second, based on the coordinates of the *i*^th^ and *i+*1^th^ parent beads in the low resolution structure, denoted as 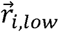 and 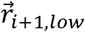, we define the coordinates of the two small derived beads from the *i*^th^ parent bead, denoted as 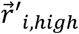 and 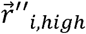, according to the following:

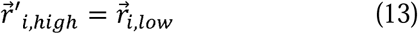

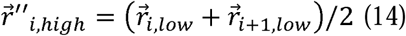

From the 3D coordinates of all the small derived beads, further 10,000 optimization steps without structure shrinking are performed to minimize the potential energy *E*(***R, C***) and output the optimized structure of high resolution. We continue the cycle until achieving the optimized structure at desired resolution. For each single cell the whole calculation is repeated 20 times based on the same experimental restraints, but starting from different random structures, thereby producing 20 structure replicas.

Here, we present the steps for the calculation of a 10-kb resolution structure as an example. First, initial whole genome structure consisting of beads representing 1280-kb regions of chromosome sequence was randomly generated and then optimized following the procedure described in the section “Maximization of conditional probability”. After that, the optimized 1280-kb resolution structure was transformed into a 640-kb resolution structure through dividing each 1280-kb bead into two 640-kb beads. Such 640-kb resolution structure was used as the initial structure for the calculation including 10,000 optimization steps without structure shrinking to output the optimized structure of 640-kb resolution. In the same manner, we obtained the optimized structures of 320 kb, 160 kb, 80 kb, 40 kb, 20 kb and final 10 kb resolution. The calculations for the 100-kb and 1-kb resolution optimized structures are starting from the initial random structures of 1600-kb and 1024-kb resolutions, respectively.

### Assigning contact read to chromatin bead pair

After dividing chromosomes into consecutive and equally sized beads, the contact read derived from the Hi-C experiment is assigned to the bead pair whose genomic positions correspond to those of the restriction fragment ends of the contact read. Therefore, we can obtain a contact map with its resolution corresponding to the size of bead. Because a hierarchical protocol is employed, calculations are performed for a series of resolutions. For instance, to obtain the 10-kb optimized structure, calculations for 1280-kb, 640-kb, 320-kb, 160-kb, 80-kb, 40-kb, 20-kb and finally 10-kb resolutions were carried out. To efficiently map Hi-C contact reads into chromatin beads of various sizes, the Si-C approach firstly divides chromosomes into 1-kb beads, and then map all contact reads into such size beads. Secondly, the Si-C approach merges consecutive, 1-kb size beads into a specified size bead to obtain a new contact map of specified resolution and simultaneously accomplish the assignment of contacts in the new map. With the increase of the size of bead, the case that the two ends of a contact share a same bead might occur. The contacts associated with such case are ignored during the structure calculations because they do not provide meaningful information on structure.

## Code and data availability

The source code of the Si-C method is available at https://github.com/TheMengLab/Si-C/tree/master/modeling. Codes for compartment identification and structural analysis are presented at https://github.com/TheMengLab/Si-C/tree/master/analysis. Calculated 3D structures by Si-C are presented at https://github.com/TheMengLab/Si-C_3D_structure. 3D structures generated by Nucdynamics are presented at https://github.com/TheMengLab/Nuc_3D_structure_Cell1. Experimental data obtained from published work has been summarized in Supplementary Note 10. Other data that support the findings of the study are available from the corresponding author upon reasonable request.

## Acknowledgements

We thank South China Normal University for financial support (Luming Meng). This work is also supported by the National Natural Science Fund of China NSFC 81502423, the Natural Science Foundation of Shanghai 20ZR1427200 and SJTU Chen Xing Type B Project 16×100080032 (Yi Shi).

## Author Contributions

Luming MENG conceived of the study, developed the Si-C method, and devised the analyses. Chenxi WANG performed the calculations and participated in analyzing the data. Yi Shi helped to interpret the experimental data downloaded from Gene Expression Omnibus (GEO) repository. Luming MENG supervised the work. Luming MENG. and Qiong Luo. wrote the paper with input from all of the authors.

