## Supplemental Materials for "Si-C: method to infer biologically valid super-resolution intact genome structure from single-cell Hi-C data"

**Table of contents**

**Supplementary Note 1. The process of calculating 3D genome structure by NucDynamics**

**Supplementary Note 2. Computing resources**

**Supplementary Note 3.** **Calculating root mean square deviation (RMSD)**

**Supplementary Note 4.** **Translating calculated 3D genome structure to distance matrix**

**Supplementary Note 5. Identifying boundaries of domain structures in distance matrix and boundaries of TADs in the populated Hi-C data**

**Supplementary Note 6. Identification of A/B compartment and features of large-scale 3D structure of the genome**

**Supplementary Note 7. Calculation of chromosome intermingling**

**Supplementary Note 8. Calculating gyration radius for chromatin region**

**Supplementary Note 9. Source list of experimental data and pre-calculation**

**Supplementary Note 1. The process of calculating 3D genome structure by NucDynamics**

To reconstruct 3D genome structure ensemble of cell 1 for the comparisons shown in Fig. 2b and 2c, we downloaded the source code of **NucDynamics** from the website <https://github.com/TheLaueLab/nuc_dynamics>. For the calculation to generate a 10-kb resolution structure ensemble including 20 conformations, we executed the **NucDynamics** software by the command as follows:

./nuc_dynamics Cell_1_contact.ncc –m 20 –f pdb –o Cell_1_10kb_20replica.pdb –s 10.24 5.12 2.56 1.28 0.64 0.32 0.16 0.08 0.04 0.02 0.01 –cpu 20

where Cell_1_contact.ncc is the Hi-C data of Cell 1 that is the exactly same data used for the structure determination of the Si-C method. Because a hierarchical protocol is employed in the NucDynamics framework, calculations were performed at 10240-kb, 5120-kb, 2560-kb, 1280-kb, 640-kb, 320-kb, 160-kb, 80-kb, 40-kb, 20-kb and finally 10-kb resolution. The values in the list “10.24 5.12 2.56 1.28 0.64 0.32 0.16 0.08 0.04 0.02 0.01” in the command represents the mentioned resolutions. For instance, the value of 10.24 means 10240-kb resolution and the last value in the list, 0.01, means the resolution of final output structure is 10 kb. “Cell_1_10kb_20replica.pdb” in the command is the name of the output file which includes 20 calculated structure replicas of 10-kb resolution. In the same manner, we generated 20-kb structure ensemble including 20 conformations by the command as follows:

./nuc_dynamics Cell_1_contact.ncc –m 20 –f pdb –o Cell_1_20kb_20replica.pdb –s 10.24 5.12 2.56 1.28 0.64 0.32 0.16 0.08 0.04 0.02 –cpu 20

, and so on for other resolution structure ensemble calculations.

**Supplementary Note 2. Computing resources**

Cpus of Intel(R) Xeon(R) CPU E5-2692 v2 @ 2.20GHz were used to compare Si-C and NucDynamics in terms of computation cost.

**Supplementary Note 3.** **Calculating root mean square deviation (RMSD)**

Before assessing the variability within the 3D genome structure ensemble calculated from sparse single-cell Hi-C data, it should be noted that the reconstructed structures are not well defined, since there are some genome regions within which no contacts were detected by the single-cell Hi-C experiments of all 8 cells. We named such regions as void regions. A brief description of the process of identifying void regions is the following. First, we divided chromosomes into beads representing 800-kb region of chromosome sequence. Second, we mapped contact reads derived from the all 8 Hi-C datasets to the beads and identified the beads where no contacts are observed as void regions. Models of 400-kb, 200-kb, and 100-kb resolutions share void regions with the 800-kb resolution model. In the same way, we identified void regions in 640-kb and 512-kb resolution models. Therefore, models of (320-kb, 160-kb, 80-kb, 40-kb, 20-kb and 10-kb) resolutions and (56-kb, 128-kb, 64-kb, 32-kb, 16-kb, 8-kb, 4-kb, 2-kb and 1-kb) resolutions share the void regions with the 640-kb and 512-kb resolution models, respectively. The void regions were excluded from the analyses of structural variability between the ensemble members.

Root mean square deviation (RMSD) is widely used to measure structural variability. In this study, we calculated RMSD between conformations within a structural ensemble according the algorithm reported by Theobald{Kim, 2019 #3416}[1], where the RMSD between two conformations is defined as:

$RMSD=\min_{trans+rot}\{\sqrt{\frac{1}{n}\sum_{i=1}^{n} \left( \vec{r}_{i,1}-\vec{r}_{i,2} \right)^{2}}\}$	(S1)

in which $\vec{r}_{i,1}$ and $\vec{r}_{i,2}$ are the coordinates of the *i^th^* bead of the two conformations, *n* is the total number of beads taken into account for the RMSD calculation. The value of RMSD is a minimum value obtained by optimally aligning the two conformations through translation and rotation.

One problem of structure reconstruction from Hi-C data is that misreconstruction such as mirror images can not be distinguished by the Hi-C experiment. Although the conformation appears in the same spatial folding as its mirror image, the RMSD between them would be high. Therefore, the variability within the ensemble including image-mirror conformations will be seriously overestimated. To overcome the issue, when calculating the pairwise RMSD between pair of conformations, denoted as Conformations (1, 2) (the numbering is quite arbitrary), we firstly constructed a image-mirror structure for Confromation 1, denoted as Conformation 1$'$ and then calculated RMSD twice, one for Conformations (1, 2), and the other for Conformations (1$'$, 2). The smaller RMSD is retained to describe the variability between Conformation (1, 2).

For convenience of the comparisons displayed in Fig. 2d, we set nuclear radius to the unity of RMSD based on the implicitly assume that intact genome 3D structure of each cell investigated here is a sphere of the same size. The nuclear radius is defined as the maximum spatial distance (in the unit of bead diameter) among the distances between every bead and the centroid of all beads.

Code for calculating RMSD is available at: <https://github.com/TheMengLab/Si-C/tree/master/analysis/structure_analysis/analysis/align/rmsd>

**Supplementary Note 4.** **Translating calculated 3D genome structure to distance matrix**

For each calculated 3D genome structure, one can measure the spatial distance between each pair of beads in the 3D structure and translate the structure into a distance matrix where matrix element represent the spatial distance between corresponding beads in the 3D genome structure. It should be noted that the value of matrix element of each distance matrix shown in Fig. 3a is computed by averaging the distance between each pair of beads across the whole 20 members of the same 10-kb structure ensemble.

Code for calculating distance matrix is available at：<https://github.com/TheMengLab/Si-C/tree/master/analysis/structure_analysis/analysis/align/distmatrix/chr1_30Mb_40Mb>/

**Supplementary Note 5. Identifying boundaries of domain structures in distance matrix and boundaries of TADs in the populated Hi-C data**

Separation score is used to quantify the degree of separating the upstream and downstream chromatin regions of one specific sequence position. The separation score is calculated from the distance matrix. Specifically, the separation score of each position is computed by averaging all the spatial distances between any pair of positions separately located in the two 500 kb regions on either side of the position. Code for Separation Score calculation is available at: <https://github.com/TheMengLab/Si-C/tree/master/analysis/structure_analysis/analysis/align/sepscore_gyr>

The positions that are identified as boundaries of domain structures in distance matrix should stratify two criteria. First, the boundary position should have higher separation score than any other positions within the 400 kb regions on either side of the position. Second, the separation score of each position within the region under investigation can be calculated and their average separation score can be obtained immediately. The boundary positon should be higher than the average separation score. The code for identifying boundary position in distance matrix is available at: <https://github.com/TheMengLab/Si-C/tree/master/analysis/structure_analysis/analysis/align/sepscore_gyr/boundary_chr>

The population Hi-C data shown in Fig. 3c and 3e is downloaded from Gene Expression Omnibus (GEO) repository with accession code GSE35156. The process for the identification of TAD boundaries in the populated Hi-C data includes the following steps:

(1)	Converting the reference genome of Hi-C data from NCBI37/mm9 to GRCm38/mm10.

(2)	All chromosome chains are divided into beads of 10-kb size and all Hi-C contact reads are assigned to pairs of beads containing the corresponding restriction fragment ends. After mapping, Hi-C data is presented in three columns, one of which lists the count of contact reads and the other two columns display the genome positions corresponding to the restriction fragment ends of contact reads.

(3) Normalizing the Hi-C data by using iterative correction and eigenvector decomposition (ICE) algorithm. The Hi-C data for individual chromosome is normalized, respectively.

(4) Converting the format of normalized Hi-C data from the form of three columns to the matrix format.

(5)	Using TopDom method[2] (version 0.0.2) to identify the TAD boundaries in each chromosome. A window size of 10 is used in the identification process.

**Supplementary Note 6. Identification of A/B compartment and features of large-scale 3D structure of the genome**

The identification of chromosome compartment is calculated following a similar algorithm described in the previous work [3]. In the calculation process, we first normalized the Hi-C contact frequency matrix through dividing each matrix element by the genome-wide average contact frequency for bin pairs at the same genomic distance. Then we calculated the correlation matrix **M**, in which the element M_ij_ describes the Pearson correlation between the *i*^th^ and *j*^th^ rows of the normalized Hi-C matrix generated in the first step. Based on the correlation matrix **M**, each chromosome was partitioned into two types of regions according to the first principal component generated by principal component analysis. Between these two types of regions, the one with higher overlap with the H3K4me3 enriched regions was defined as compartment A and the other one was defined as compartment B.

The script is available at: <https://github.com/TheMengLab/Si-C/tree/master/analysis/compartment>

Supplementary Fig.1 displays several features of 3D genome architecture for Cells 2-8. Supplementary Fig.2 shows the locations of centromeres and telomeres in the nucleus for the all eight cells.

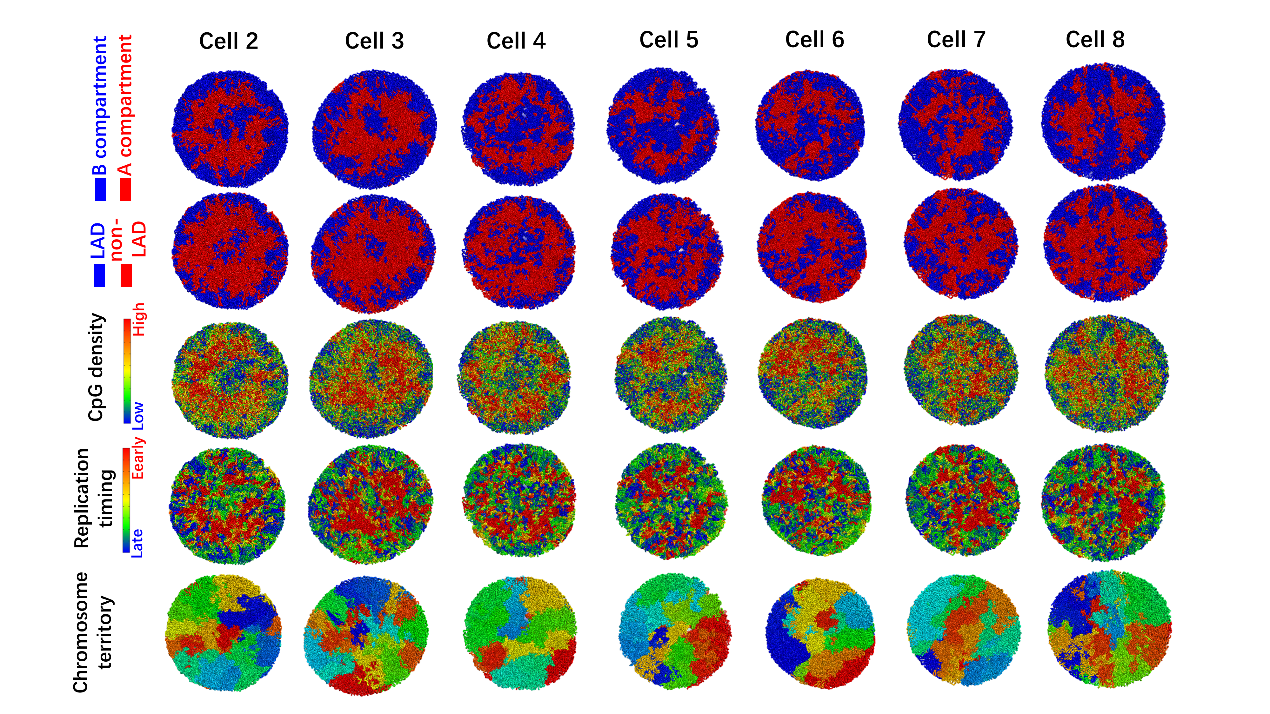

**Supplementary Figure 1.** **Cross-sections of the Si-C intact genome 3D 10-kb resolution structures of Cell 2-8.** Colored according to whether the sequence is in the A (red) or B (blue) compartment (first column); whether the sequence is part of a lamina associated domain (LAD) (blue) or not (red) (second column); the CpG density from red to blue (high to low) (third column); the replication time in the DNA duplication process from red to blue (early to late) (fourth column).

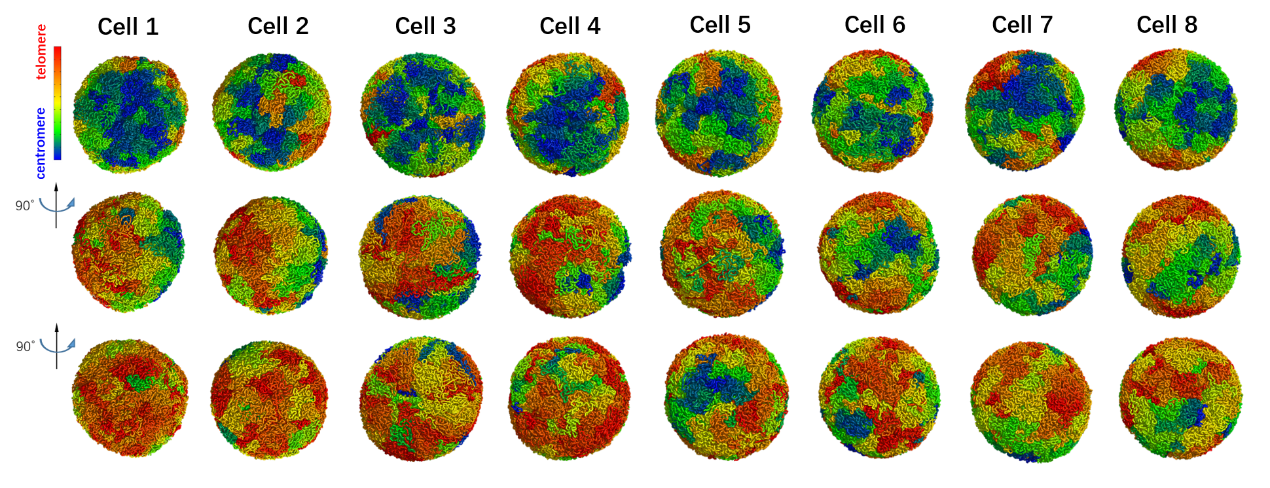

**Supplementary Figure 2. The locations of centromeres and telomeres in the Si-C intact genome 3D 10-kb resolution structures for the all eight individual cells.** A consistent Rabl configuration (with centromeres and telomeres clustered on opposite sides of the nucleus) are shown in all G1-phase ES cells, strongly validating the Si-C 10-kb resolution structures.

**Supplementary Note 7. Calculation of chromosome intermingling**

To assess the degree of intermingling between chromosomes, we first identified intermingled beads within each chromosome and then calculated the proportion of intermingled beads to the total beads of the chromosome. The intermingled beads were defined as those that surrounded by at least four other beads from a different chromosome within a distance threshold between beads of 2 bead diameters.

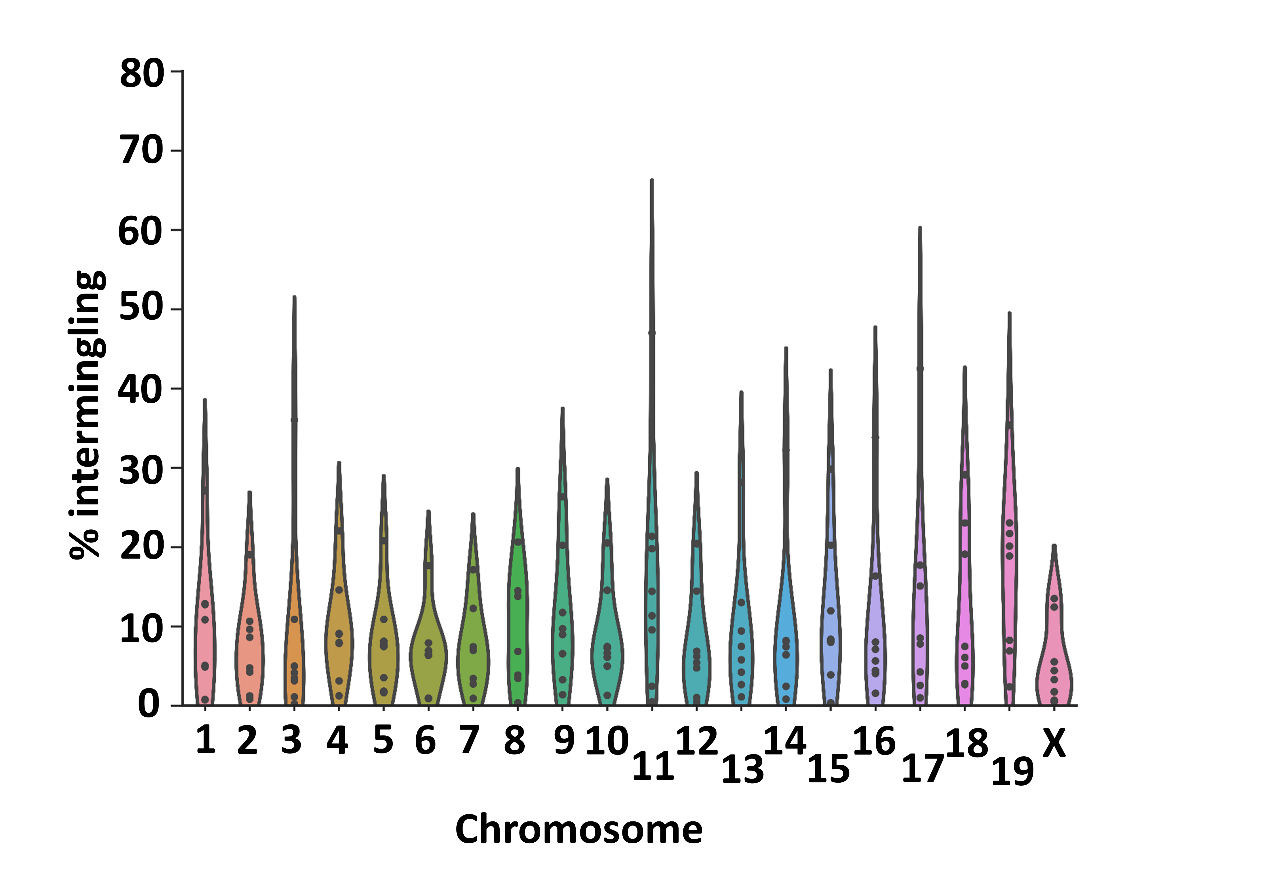

**Supplementary Figure 3.** Violin plot showing the proportion of each chromosome that intermingles with other chromosomes. The proportion is derived from the Si-C 10-kb resolution structures of the eight G1-phase ES cells,.

**Supplementary Note 8. Calculating gyration radius for chromatin region**

We estimated the degree of compaction of investigated regions of 200 kb using the gyration radius (*R_g_*) which is defined as the following:

$R_{g}=\sqrt{\frac{1}{N}\sum_{i=1}^{N} \left( \vec{r}_{i}-\vec{r}_{\mathrm{ave}} \right)^{2}}$	(S2)

in which *N* represents the number of beads in the 200 kb region under investigation, $\vec{r}_{i}$ the coordinate of the *i^th^* bead in the region and $\vec{r}_{ave}$ is the coordinate of the centroid of the region. In this study, the value of N is 20 because the region of 200 kb is represented by beads of 10-kb size.

The script for calculating is gyration radius available at: <https://github.com/TheMengLab/Si-C/tree/master/analysis/structure_analysis/analysis/align/sepscore_gyr>

**Supplementary Note 9. Source list of experimental data and pre-calculation**

The experimental data used in our analysis was taken from previously published work, as elaborated below:

| Data type | Accession number | Reference |
| --- | --- | --- |
| H3K4me3 ChIP-seq (haploid) | GSE80280 | Stevens, T.J. et al. 3D structures of individual mammalian genomes studied by single-cell Hi-C. *Nature* **544**, 59-+ (2017). |
| Constitutive Lamina Associated Domain | GSE17051 | Peric-Hupkes, D. et al. Molecular Maps of the Reorganization of Genome-Nuclear Lamina Interactions during Differentiation. *Mol Cell* **38**, 603-613 (2010). |
| Replication Timing | E-MTAB-3506 | Kolesnikov, N. et al. ArrayExpress update-simplifying data submissions. *Nucleic Acids Res* **43**, D1113-D1116 (2015). |
| Populated Hi-C data | GSE35156 | Dixon, J.R. et al. Topological domains in mammalian genomes identified by analysis of chromatin interactions. *Nature* **485**, 376-380 (2012). |
